# Complement therapeutic Factor H-IgG proteins as pre-exposure prophylaxes against Lyme borreliae infections

**DOI:** 10.1101/2024.09.26.615144

**Authors:** Connor W. McKaig, Jill Malfetano, Y Tran, Xiuli Yang, Utpal Pal, Keith Wycoff, Yi-Pin Lin

**Affiliations:** Department of Infectious Disease and Global Health, Cummings School of Veterinary Medicine, Tufts University, North Grafton, MA, USA; Division of Infectious Diseases, Wadsworth Center, NYSDOH, Albany, NY, USA; Planet Biotechnology, Inc., Hayward, CA, USA; Department of Veterinary Medicine, Virginia-Maryland Regional College of Veterinary Medicine, University of Maryland, College Park, MD, United States

**Keywords:** FH-Fc, Lyme disease, *Borrelia*, Pre-exposure prophylaxis, complement

## Abstract

Lyme disease (LD) is the most common vector-borne disease in the northern hemisphere and is caused by the bacteria *Borrelia burgdorferi* sensu lato (also known as Lyme borreliae) with no effective prevention available. Lyme borreliae evade complement killing, a critical arm of host immune defense, by producing outer surface proteins that bind to a host complement inhibitor, factor H (FH). These outer surface proteins include CspA and CspZ, which bind to the 6^th^ and 7^th^ short consensus repeats of FH (SCR(6-7)), and the OspE family of proteins (OspE), which bind to the 19^th^ and 20^th^ SCR (SCR19-20). In this study, we produced two chimeric proteins, FH-Fc, containing the Fc region of immunoglobulin G (Fc) with SCR(6-7) or SCR(19-20). We found that both FH-Fc constructs killed *B. burgdorferi* in the presence of complement and reduced bacterial colonization and LD-associated joint inflammation *in vivo*. While SCR(6-7)-Fc displayed Lyme borreliae species-specific bacterial killing, SCR(19-20)-Fc versatilely eradicated all tested bacterial species/strains. This correlated with SCR(6-7)-Fc binding to select variants of CspA and CspZ, but SCR(19-20)-Fc binding to all tested OspE variants. Overall, we demonstrated the concept of using FH-Fc constructs to kill Lyme borreliae and defined underlying mechanisms, highlighting the potential of FH-Fc as a pre-exposure prophylaxis against LD infection.

**AUTHOR SUMMARY:** Transmitted by ticks, Lyme disease (LD) is the most common vector-borne disease in North America and has experienced an expanded geographical range and increasing number of cases in recent years. No effective prevention is currently available. The causative agent of LD, *Borrelia burgdorferi* sensu lato (*Bb*sl), is a complex containing a variety of species. To escape from killing by complement, one of the mammalian host defense mechanisms, *Bb*sl produces outer surface proteins that bind to a complement inhibitor, factor H (FH). These FH-binding proteins (i.e., CspA, CspZ, and OspE) evade complement by recruiting FH to the bacterial surface. Here we produced two FH-Fc fusion proteins, which combine human immunoglobulin Fc with the human FH domains that bind to *Bb*sl FH-binding proteins. We found that FH-Fc constructs kill *Bb*sl *in vitro* and prevent colonization and LD manifestations in murine models, correlating with these FH-Fc constructs’ ability to bind to CspA, CspZ, and OspE from respective *Bb*sl species. These results suggest the possibility of using FH-Fc as a prevention against LD.

## INTRODUCTION

Lyme disease is the most common vector-borne disease in the northern hemisphere, and the disease incidence is escalating: the CDC estimated more than 476,000 cases in the United States and approximately 10,000 cases are reported each year in Europe (1–3). Transmitted by *Ixodes* ticks, Lyme disease is caused by more than 21 species of spirochete bacteria, collectively named *Borrelia burgdorferi* sensu lato (also known as *Borreliella* burgdorferi, *B. burgdorferi* s.l., or Lyme borreliae)(4). Humans can be infected by selected *B. burgdorferi* s.l. species, including *B. burgdorferi* sensu stricto (hereafter *B. burgdorferi*) prevalent in both North America and Eurasia, and *B. afzelii*, *B. bavariensis*, and *B. garinii* isolated from Eurasia (5). Each of these species have evolved into multiple genetically distinct “strains”, which differ in their associated manifestations and the incidence of human cases (5). Following a tick bite, the bacteria disseminate through the bloodstream from the bite site on the skin to multiple tissues and organs, causing manifestations, such as arthritis, neuroborreliosis, carditis, and acrodermatitis(6–10). Despite the continuing rising cases, geographical expansion of prevalence, and diversity of causative agents, no effective LD preventive is currently available.

The survival of Lyme borreliae in humans and reservoir animals requires the ability to overcome different arms of the vertebrate host immune response in the blood, and the first-line defense is the complement system (11–13). Complement can be activated through three canonical pathways: the classical and lectin pathways are initiated by the localization of antigen-antibody complexes and mannose-binding lectin (MBL) on the surface of microbes, respectively; the alternative pathway is initiated by C3b binding to the microbial surface (**Fig. 1A**) (14). Activation results in the formation of either of two C3 convertase enzymatic complexes: (1) C4b2a, whose assembly is triggered via the classical or lectin pathways; and (2) C3bBb, enzymatic complexes, triggered via the alternative pathway **(Fig. 1A)**. Both C3 convertases induce the release of proinflammatory peptides (C3a, C5a), the deposition of opsonins (iC3b) on the microbial surface, and, by recruiting other complement proteins, generate C5 convertases. The latter catalyzes the formation of the membrane attack complex, C5b-9, for pathogen lysis **(Fig. 1A)**. The proinflammatory nature of the complement cascade necessitates tight control of this potentially destructive immune defense. Indeed, the host encodes regulators of complement activation (RCA), proteins that modulate each of the three activation pathways (15). For example, factor H (FH), which contains 20 short consensus repeats (SCRs), binds to and triggers degradation of C3b to inhibit C3b-containing convertases generated by the alternative pathway(16, 17) **(Fig. 1A)**.

**Figure 1.**
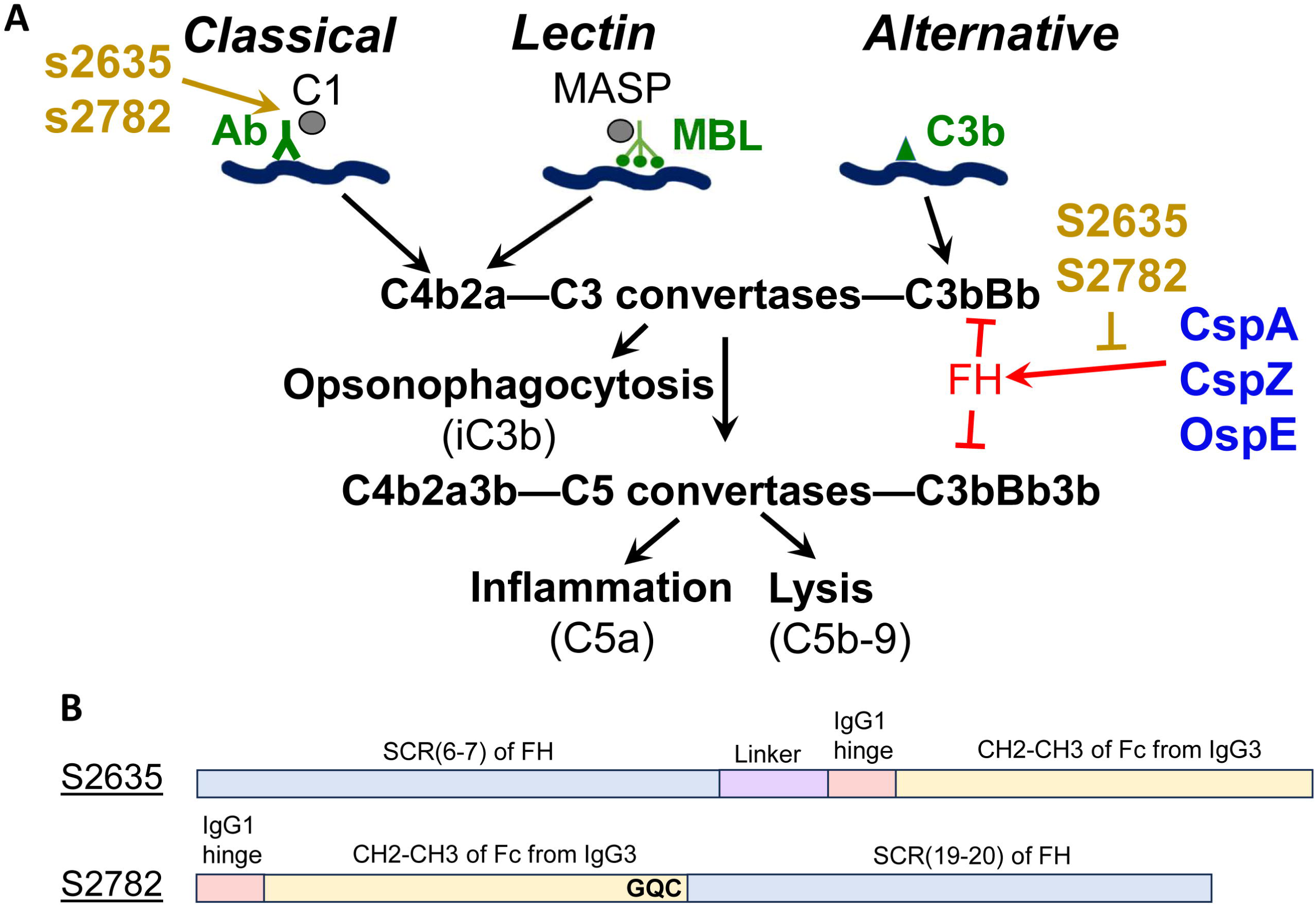
Schematic diagram showing composition of the FH-Fc constructs used in this study and their proposed mechanisms of action to eliminate Lyme borreliae. **(A)** FH-Fc is proposed to promote *B. burgdorferi* s.l. killing by binding to the FH-binding proteins of Lyme borreliae via the FH region to prevent the pathogens from escaping alternative complement pathway-mediated pathogen killing. Simultaneously, the Fc region of FH-Fc may recruit the complement C1 components (C1) required for classical complement pathway-mediated pathogen killing. Lyme borreliae produce CspA, CspZ, and OspE to bind to FH in promoting alternative pathway evasion. Shown are two FH-Fc constructs, S2635 and S2782, in binding to CspA, CspZ, and/or OspE to prevent bacterial evasion to alternative pathway and interacting with C1 to promote classical pathway activation. This panel was adopted from our previous publication (82). **(B)** Shown are the diagrams for the composition of S2635 and S2782: S2635 contains the SCR(6-7) region of human FH, a linker, the hinge region of the human IgG1, and the CH2-CH3 region of the Fc region from human IgG3 in sequential order from N to C terminus. S2782 contains the hinge region of the human IgG1, the CH2-CH3 region of the Fc region from human IgG3, and the SCR(6-7) region of human FH in sequential order from N to C terminus. The mutated amino acids at the C-terminal Fc region have been highlighted in bold. The amino acid sequences of S2635 and 2782 are indicated in **Text S1** and **S2**, respectively.

To survive in the blood where complement is mainly located, Lyme borreliae produce RCA-binding proteins to bind and recruit complement regulatory proteins on the spirochete surface to inactivate complement (12, 18). One group of these proteins bind to FH(19). Among these FH-binding proteins, CspA (also known as <u>C</u>omplement <u>R</u>egulator <u>A</u>cquiring <u>S</u>urface <u>P</u>rotein 1 or CRASP-1) and CspZ (also known as CRASP-2) bind to SCR(6-7) of FH(20, 21). The other proteins, belonging to a OspE protein family (also known as CRASP3-5), bind to SCR(19-20) of FH (20, 22, 23). During the enzootic cycle, CspA is expressed mostly when bacteria are in ticks prior to and during tick-to-vertebrate host transmission, whereas CspZ and OspE are largely produced while bacteria are in the vertebrate hosts after transmission (24). While the role of OspE in the enzootic cycle remains undefined, CspA and CspZ confer bacterial evasion to complement in ticks’ bloodmeal to facilitate tick-to-host transmission and in the host bloodstream for efficient dissemination, respectively (25–27). These findings thus led to the concept that targeting these FH-binding functions might serve as an intervention against Lyme disease.

In fact, efforts have been made to test this concept by generating several candidates of complement-targeted therapeutics, one of which is FH-Fc (28). FH-Fc are recombinant fusion proteins containing SCR(6-7) or SCR(19-20) that bind to the FH-binding proteins from pathogens, as most pathogens’ FH-binding proteins target those FH regions. By displacing FH on the pathogen surface, FH-Fc are intended to prevent alternative complement pathway evasion. FH-Fc also contain the Fc region of immunoglobulins, allowing the activation of classical pathway-mediated pathogen killing(28). In support of this concept, FH-Fc has been demonstrated to efficiently eliminate multiple bacterial or parasite species, such as *Neisseria gonorrhoeae, Neisseria meningitidis, Streptococcus pyogenes, Haemophilus influenza, Trypanosoma cruzi*(29–31, 32, 33).

In this study, we produced two FH-Fc constructs using *Nicotiana benthamiana* (tobacco plant expression system), a platform that can rapidly produce large amounts of foreign proteins for pharmaceutical use (34). We tested the ability of these FH-Fc constructs to kill different species or strains of *B. burgdorferi* s.l. *in vitro* and to prevent Lyme-associated bacterial colonization and manifestations using murine models. We also attempted to determine the mechanisms underlying the FH-Fc-mediated Lyme borreliae killing by measuring the binding affinity of FH-Fc constructs to bacterial FH-binding proteins to investigate the potential of FH-Fc as a Lyme disease prophylaxis.

## RESULTS

### S2635 and S2782 were constructed and produced in *Nicotiana benthamiana*

To generate the SCR6-7 and SCR19-20 versions of FH-Fc constructs (S2635 and S2782, respectively), we obtained the plant codon-optimized DNA of those FH domains. The SCR6-7 sequence in S2635 was then connected at the N-terminal end of the CH2-CH3 domains (Fc) of human IgG3, with SCR6-7 and Fc separated by a flexible linker (GGGGSGGGGSGGGGSS), followed by a portion of the IgG1 hinge sequence (EPKSCDKTHTCPPCP) (**Fig. 1B**, **Text S1**). The N-terminus of S2782 starts with a portion of IgG1 hinge sequence (DKTHTCPPCP) and the human CH2-CH3 domains from IgG3, with a small change in which the C-terminal three residues, PGK, were replaced by the GQC (**Fig. 1B**, **Text S2**). Such replacement was intended to facilitate the resistance to proteolytic cleavage between Fc and SCR19-20 of FH (35). Additionally, adding a cysteine was intended to promote the formation of an inter-chain disulfide bond between paired CH3 domains to stabilize the construct against aggregation by low pH (35). The plant codon-optimized DNA sequence encoding human SCR19-20 was then appended to the C-terminal end of the IgG3 Fc (**Fig. 1B**, **Text S2**). Both S2635 and S2782 were then produced using a rapid *N. benthamiana* expression system and purified by protein A affinity chromatography (31). The resulting yields of purified S2635 and S2782 were 296 ±23 and 522 ±172 mg/kg, respectively.

### S2635 and S2782 differ in their ability to eliminate *B. burgdorferi* in fed nymphs, but both significantly reduced bacterial colonization and joint inflammation

We tested the ability of S2635 or S2782 to impact bacterial colonization and Lyme disease-associated manifestations in mice. One day prior to nymphal tick feeding (−1 dpf), we injected mice with 0.2 mg/kg of S2635, S2782 or PBS (control) intramuscularly, as immunoglobulins and Fc-fusions are highly bioavailable when introduced by this route (36)) (**Fig. 2A**). At 24-h after the injection, we allowed *I. scapularis* nymphal ticks carrying *B. burgdorferi* strain B31-5A4 to feed on the mice. Uninfected mice (PBS-pre-treated mice without tick feeding) were also included as a control (**Fig. 2A**). We first measured the bacterial burdens in replete nymphs after engorgement and found that the nymphs feeding on S2635- but not S2782-pre-treated mice had significantly lower burdens than those feeding on PBS-pre-treated mice. These results suggest the ability of S2635 to uniquely eliminate the bacteria in feeding ticks during tick-to-host transmission (**Fig. 2B**).

**Figure 2.**
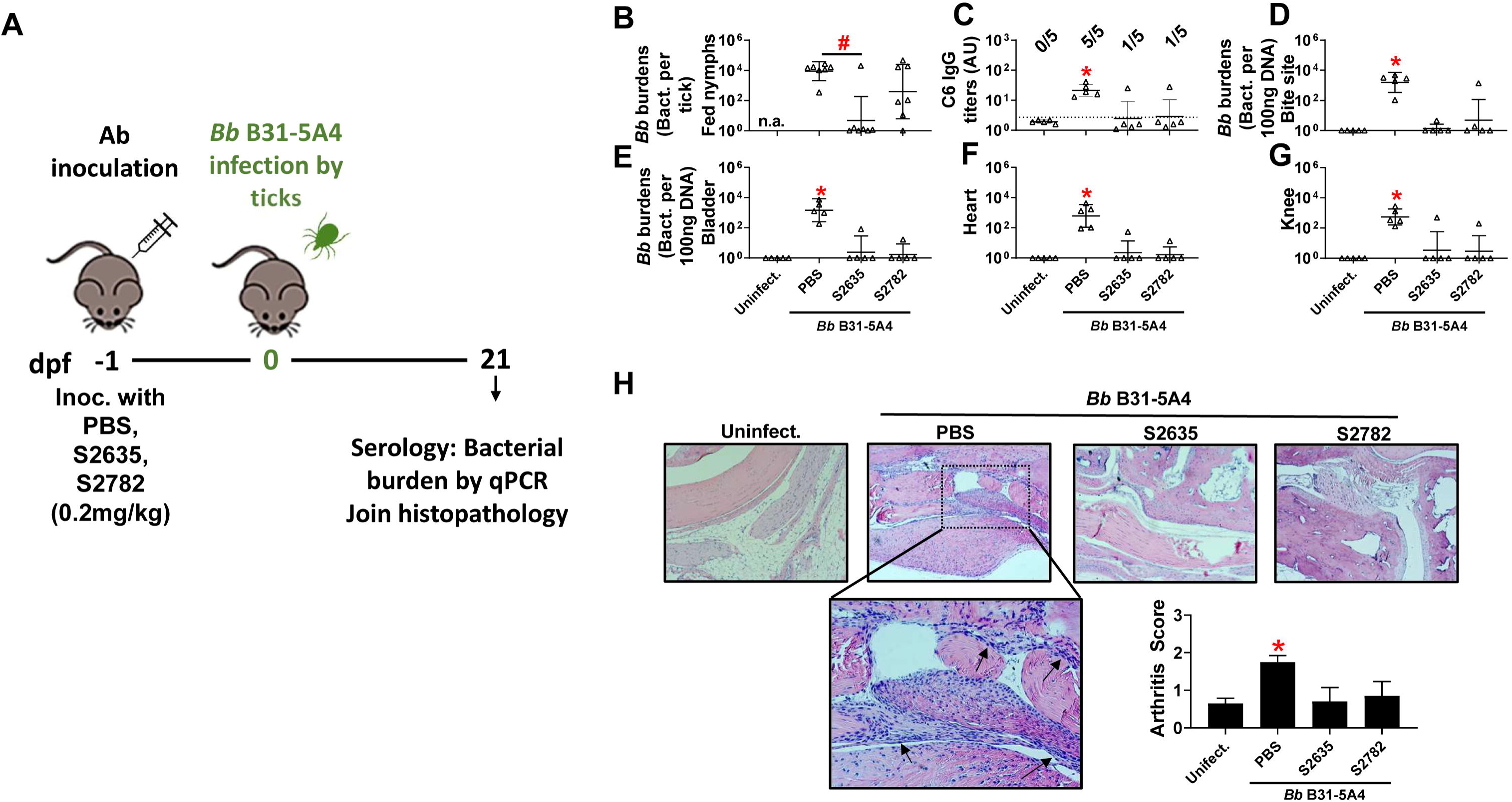
S2635 and S2782 protected mice from Lyme borreliae-associated colonization, seropositivity, and arthritis but differed in the ability to eliminate spirochetes in fed nymphs. **(A)** Shown is the timeframe of FH-Fc inoculation and Lyme borreliae infection in this study. **(B to H)** Five C3H/HeN mice were intramuscularly injected with S2635 or S2782 at the dose of 0.2 mg/kg, or PBS (control). At 24 hours after inoculation (1-day prior to tick feeding (dpf)), these mice were fed on by *I. scapularis* nymphs carrying *B. burgdorferi* B31-5A4 (*Bb* B31-5A4). An additional five mice injected with PBS but not fed on by ticks were included as the control (Uninfect.). **(B)** The engorged fed nymphs were collected from those mice at 4 dpf. **(C)** At 21 days, sera were collected from these mice to determine the seropositivity to Lyme disease infection by evaluating the IgG levels of C6 antigens. Spirochete burdens at **(D)** the tick feeding site (“Bite Site”), **(E)** bladder, **(F)** heart, and **(G)** knees were quantitatively measured at 21 dpf, shown as the number of spirochetes per 100ng total DNA. Data shown are the geometric mean ± geometric standard deviation of the spirochete burdens from five mice per group. Statistical significances (p < 0.05, Kruskal-Wallis test with the two-stage step-up method of Benjamini, Krieger, and Yekutieli) of differences in bacterial burdens relative to (*) uninfected mice are presented. **(H)** Tibiotarsus joints at 21dpf were collected to assess inflammation by staining these tissues using hematoxylin and eosin. Representative images from one mouse per group are shown. Top panels are lower-resolution images (joint, ×10 [bar, 160 µm]); bottom panels are higher-resolution images (joint, 2×20 [bar, 80 µm]) of selected areas (highlighted in top panels). Arrows indicate infiltration of immune cells. **(Inset figure)** To quantitate inflammation of joint tissues, at least ten random sections of tibiotarsus joints from each mouse were scored on a scale of 0-3 for the severity of arthritis. Data shown are the mean inflammation score ± standard deviation of the arthritis scores from each group of mice. Asterisks indicate the statistical significance (p < 0.05, Kruskal Wallis test with the two-stage step-up method of Benjamini, Krieger, and Yekutieli) of differences in inflammation relative to uninfected mice.

At 21-dpf, we measured the spirochete burdens in mouse tissues and Lyme borreliae seropositivity (i.e., the IgG levels against a C6 peptide derived from a Lyme borreliae VlsE antigen, a commonly used biomarker for Lyme disease serodiagnosis (37)) (**Fig. 2A**). PBS-pre-treated mice yielded significantly greater levels of seropositivity (five out of five turning seropositive) than uninfected mice (**Fig. 2C**). In contrast, only one out of five S2635- or S2782-pre-treated mice turned Lyme borreliae seropositive, which is statistically indistinguishable from uninfected mice (**Fig. 2C**). Additionally, compared to uninfected mice, all 5 PBS-pre-treated mice had significantly higher bacterial burdens at tick bite sites (skin) and in bladder, heart, and knee joints, whereas the same tissues from only 1 out of 5 mice pre-treated with S2635 or S2782 had detectable bacterial burdens (**Fig. 2D to G**). These results indicate the ability of S2635 and S2782 to reduce bacterial colonization after *B. burgdorferi* tick-to-host transmission.

We further determined the Lyme disease-associated manifestations in those mice at 21-dpf by histological analysis of the ankle joints. In PBS-pre-treated mice, we observed elevated levels of infiltrations of granulocytes and mononuclear cells, such as neutrophils and monocytes in the connective tissues, tendons, and muscles (**Fig. 2H**). That resulted in significantly greater levels of inflammation scores in PBS-pre-treated mice than those in uninfected mice (**Fig. 2H**, inset figure). However, we did not observe such noticeable cell infiltrations in the joints of S2635- and S2782-pre-treated mice, agreeing with their indistinguishable inflammatory scores, compared to the scores of uninfected mice (**Fig. 2H**). Overall, these findings indicated that S2635 or S2782 pre-treatment prior to nymph transmission of *B. burgdorferi* B31-5A4 efficiently decreased spirochete colonization and Lyme disease-associated joint inflammation.

### *In vitro* complement-directed killing of Lyme borreliae by S2635 and S2782

We next examined the ability of S2635 and S2782 to kill *B. burgdorferi* B31-5A4 *in vitro*. Human serum (the source of complement) was incubated with strain B31-5A4 in the presence of different concentrations of S2635, S2782, or BSA (control). FH-Fc constructs are proposed to eradicate spirochetes by promoting classical pathway-mediated killing via their Fc regions and preventing alternative pathway evasion by displacing FH from the bacteria. Therefore, 40% human serum was used as the final concentration in the reaction because this percentage allows for the observation of Lyme borreliae killing by both pathways. However, *B. garinii* strains ZQ1 and PBr were found to be incapable of surviving this concentration of human serum (**Fig. S1**). We thus titrated the serum and found 20% as the minimal concentration that these two strains can survive and thus incubated these strains in 20% human serum for this experiment (**Fig. S1**). We then quantified the bacteria present microscopically after 4-h of incubation and then normalized those numbers to those prior to the incubations. Such normalization permitted us to measure the percent survival of bacteria, calculating the EC_50_ values (the concentration of FH-Fc constructs that lead to 50% of bacterial survival). While BSA-incubated *B. burgdorferi* yielded close to 100% of bacterial survivability, S2635 and S2782 incubations resulted in spirochete killing, with S2635 having significantly more robust killing than S2782 based on their EC_50_ values (**Fig. 3A and Table S1**).

**Figure 3.**
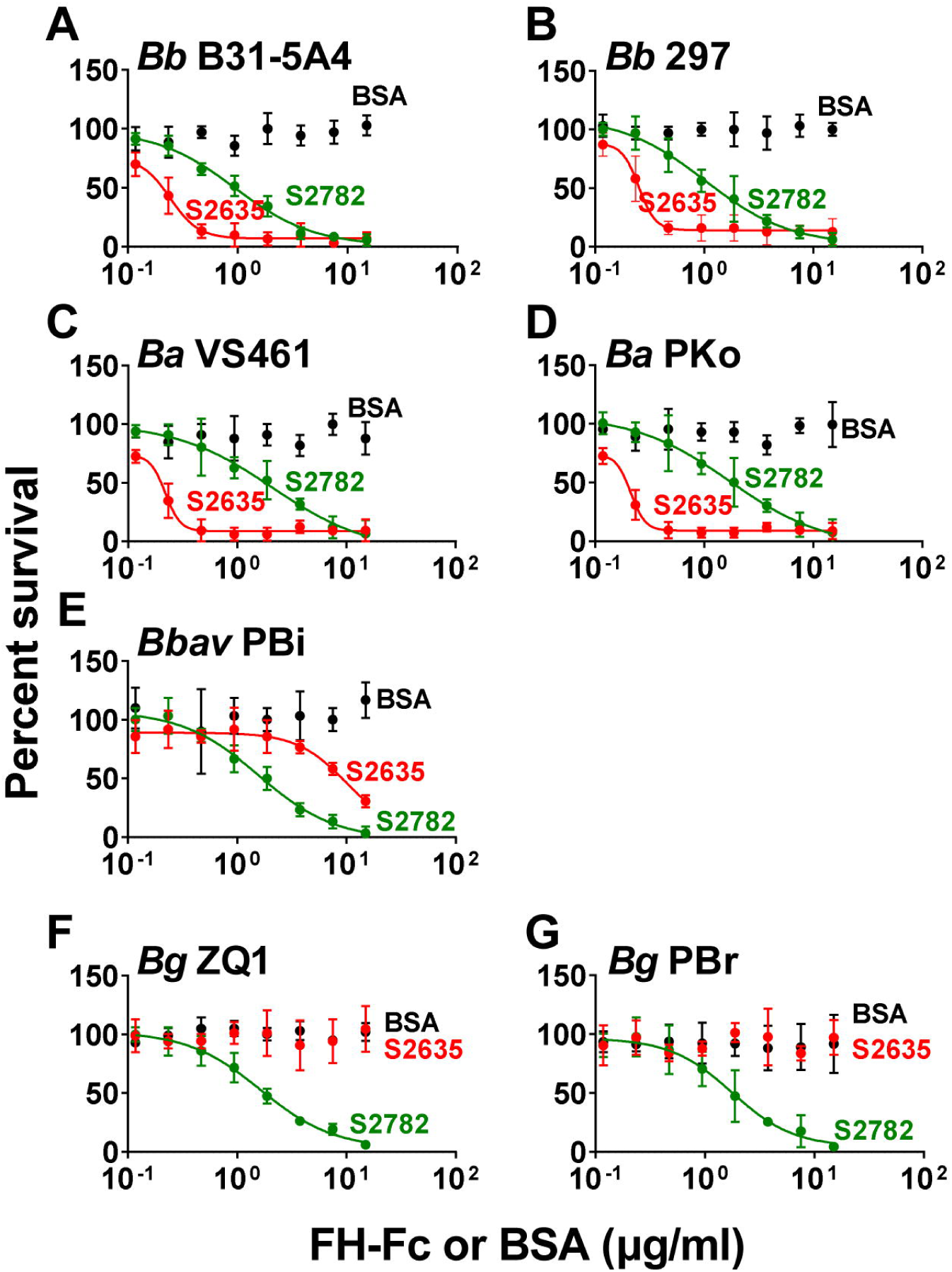
S2635 and S2782 varied in their breadth of killing to different Lyme borreliae species and strains. S2635, S2782, or BSA (control) or PBS (control, data not shown) were serially diluted as indicated, and mixed with human serum and incubated with the following Lyme borreliae species and strains (5 × 10^5^ cells ml^−1^): *B. burgdorferi* (*Bb*) strains **(A)** B31-5A4 and **(B)** 297, *B. afzelii* (*Ba*) strains **(C)** VS461 and **(D)** PKo, *B. barvariensis* (*Bbav*) strain **(E)** PBi, and *B. garinii* (*Bg*) strains **(F)** ZQ1 and **(G)** PBr. The final concentration of the human serum was 40% except that of *B. garinii* strains which was 20%. After incubating for 24 hours, surviving spirochetes were quantified from three fields of view for each sample using dark-field microscopy. The work was performed on three independent experiments. The survival percentage was derived from the proportion of FH-Fc-treated to PBS-treated spirochetes. Shown is one representative experiment, and in that experiment, the data points are the mean ± SEM of the survival percentage from three replicates. The 50% borreliacidal activity of each FH-Fc (EC_50_), representing the FH-Fc concentrations that effectively killed 50% of spirochetes, was obtained and extrapolated from curve-fitting and shown in **Table S1**. The EC_50_ values are shown as the mean ± SD of from three experiments.

We next examined whether such bactericidal activity of S2635 and S2782 can be extended to other Lyme borreliae species and strains and obtained EC_50_ values for those bacteria in the same fashion. Our results showed three patterns of bacterial killing depending on the Lyme borreliae species or strains mixed with S2635 or S2782: 1) Similar to *B. burgdorferi* B31-5A4, when incubated with *B. afzelii* strains VS461 and PKo, both S2635 and S2782 efficiently killed those strains. However, S2635 eradicated these strains more efficiently than S2782 (**Fig. 3B to D and Table S1**); 2) When incubated with *B. bavariensis* strain PBi, although both S2635 and S2782 can eliminate this strain, S2782 showed more significantly robust killing than S2635 (**Fig. 3E and Table S1**); 3) When incubated with *B. garinii* strains ZQ1 and PBr, S2635 did not show bacterial killing, but S2782 was found to kill those strains (**Fig. 3F and G**). These results support the versatility of S2782-mediated spirochete killing but unique strain-specific Lyme borreliae killing by S2635.

### Binding affinity of S2635 and S2782 to purified Lyme borreliae FH-binding proteins

We hypothesized that the *in vitro* killing potency of these two FH-Fc would depend on their affinity for pathogen FH-binding proteins. To gain insights into the molecular basis of FH-Fc-mediated Lyme borreliae killing, we produced recombinant forms of several Lyme borreliae FH-binding proteins, including CspA, CspZ, and OspE, from four different species (19). We produced one variant per FH-binding protein from each of the tested species based on sequences that are available in GenBank, with the exception that variants from two strains of *B. burgdorferi* (B31 and 297) were produced. Note that two CspA paralogs from *B. bavariensis* (Bga66 and Bga71) (38) and two OspE paralogs from *B. burgdorferi* strain B31 (ErpA and ErpP) have been shown to bind human FH (39, 40). Therefore, those variants were also included in the study. S2635 and S2782 were conjugated to separate SPR chips, which were used to measure binding affinity of each of the Lyme borreliae FH-binding proteins. We did not detect binding of any of the OspE variants to S2635 (**Fig. 4A, top panel, Table S2)**, in agreement with prior findings that OspE does not bind to human SCR6-7 (41). However, we found that S2635 binds strongly to the CspA variants from *B. burgdorferi* strains B31 and 297 and *B afzelii* strain PKo (K_D_ = 1.1 to 2.2×10^−7^ M, **Fig. 4B top panel, Table S3**) but less efficiently to *B. bavariensis* strain PBi (K_D_ = 1.1×10^−6^ M, **Fig. 4B top panel, Table S3**). No binding of S2635 to the CspA variant of *B. garinii* strain ZQ1 was detected (**Fig. 4B top panel, Table S3**). Similarly, binding of CspZ to S2635 was species- and strain-specific: S2635 bound robustly to the CspZ variants from *B. burgdorferi* strain B31 and *B. afzelii* (K_D_ = 1.5 to 2×10^−7^ M, **Fig. 4C top panel, Table S4**), but less efficiently to the CspZ variant from *B. bavariensis* (K_D_ = 9.8×10^−7^ M, **Fig. 4C top panel, Table S4**). Additionally, S2635 did not bind to the CspZ variants from *B. burgdorferi* strain 297 or *B. garinii* (**Fig. 4C top panel, Table S4**). We also applied each of these CspA, CspZ, and OspE variants to the S2782-conjugated SPR chips. S2782 did not bind to any tested CspA or CspZ variants (**Fig. 4B and C bottom panels, Table S3 and S5**), consistent with the inability of these FH-binding proteins to bind to human SCR19-20 (21, 42). However, we detected the binding of all tested OspE variants to S2782, indicating the versatility of the S2782-binding ability of OspE (**Fig. 4A bottom panel, Table S2**).

**Figure 4.**
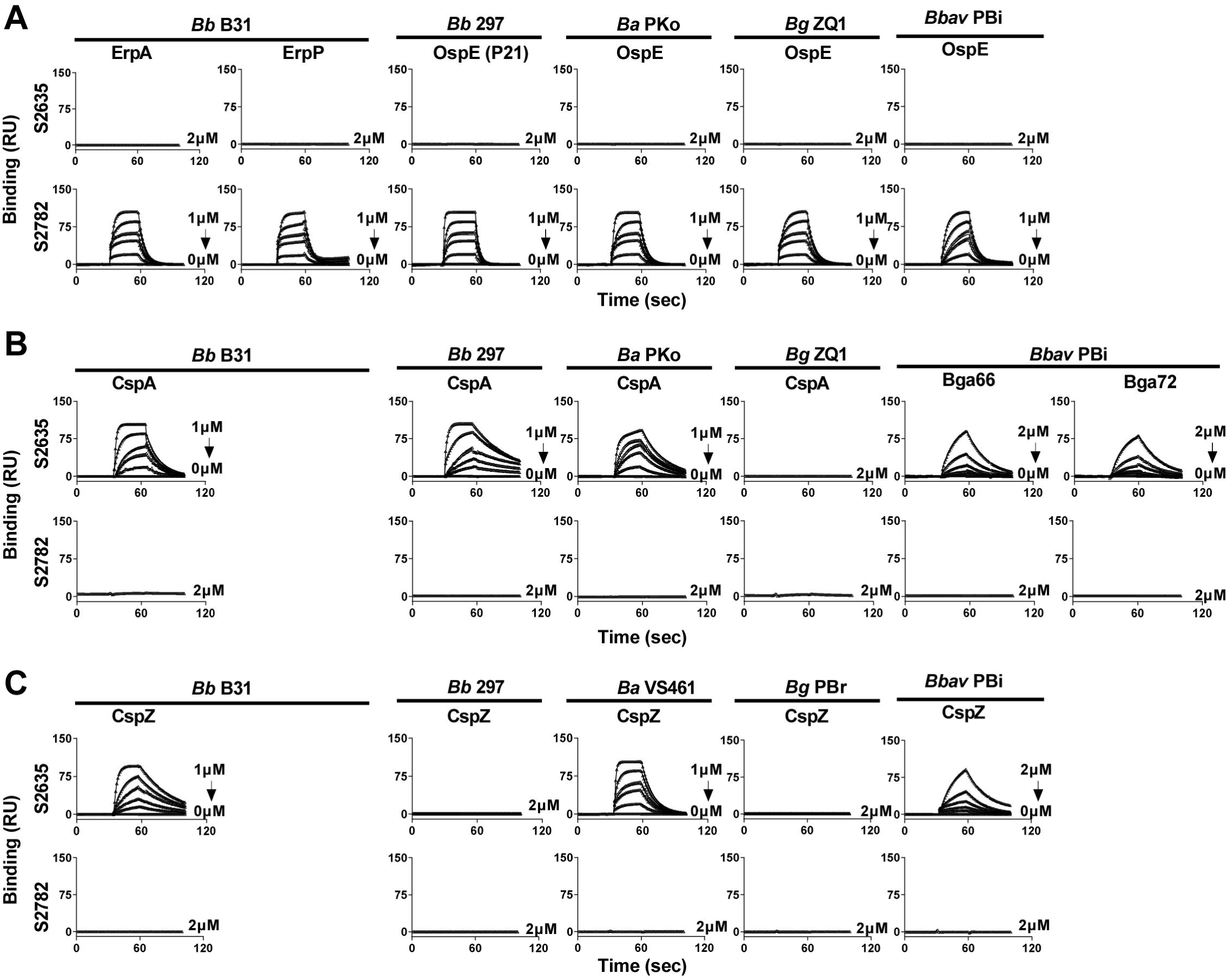
S2635 and S2782 bound to Lyme borreliae factor H-binding protein variants in a variant-specific manner. The indicated concentrations of the variants of **(A)** OspE, **(B)** CspA, or **(C)** CspZ from indicated strains *B. burgdorferi* (*Bb*), *B. afzelii* (*Ba*), *B. garinii* (*Bg*), or *B. bavariensis* (*Bbav*) were flowed in PBS buffer over the chip surface, conjugated with S2635 or S2782. Binding was measured in response units (R.U.) by SPR. The k_on_, k_off_, and K_D_ values were determined from the average of three experiments **(Table S2, S3, and S4)**. Shown is one representative experiment.

## DISCUSSION

As a one of the major host defense mechanisms, complement not only lyses pathogens but cross talks with different arms of the host immune response (43–45). Dysregulation of this defense mechanism thus often causes various autoimmune diseases and exacerbates infectious diseases. Manipulating complement regulation by targeting complement components or regulators has frequently been employed in developing therapies for those autoimmune and infectious diseases (46). Specifically, to combat complement dysregulation-mediated infectious diseases, one of the most commonly used strategies is to target pathogens’ anti-complement proteins either by monoclonal antibodies or small molecules (47). However, targeting single pathogen proteins may risk pathogens developing “neutralization escape mutants”, leading to ineffective therapeutics (48). Additionally, the functional redundancy seen in pathogen anti-complement mechanisms makes this a complex strategy(49). An alternative approach is to target the host complement components or regulators that are involved in pathogen complement evasion to skew the complement responses toward pathogen elimination (50). Such therapeutics would be mimetics of the pathogen-binding domains of host complement regulators, preventing pathogens from hijacking host complement or eliciting complement regulators to inhibit complement activation (50). Some of these “complement-based” therapeutics have been produced as fusion proteins with the Fc of immunoglobulin, such as FH-Fc to further enhance classical pathway-mediated pathogen killing (29–32). It should be more difficult for pathogens to evolve neutralization escape mutations to FH-Fc, as any mutations that reduce FH-Fc binding should have reduced FH binding, leading to more robust complement-mediated killing (29–32, 51). Moreover, as binding to complement regulator is a common approach shared by many pathogens to evade complement, complement modulation-targeted therapeutics (e.g., FH-Fc) have the potential to be applied as broad spectrum anti-infectives.

In this study, both FH-Fc constructs (i.e., S2635 and S2782) efficiently killed *B. burgdorferi in vitro*, and reduced *B. burgdorferi* dissemination and the development of Lyme disease-associated manifestations in the murine model when administered prior to tick challenge. However, pre-treatment of mice with S2635 but not S2782 eliminated bacterial burdens in fed nymphs, suggesting that the bacterial killing mechanisms for these FH-Fc constructs differ. One contributor of such differences could be the infection stages when the bacterial FH-binding proteins are produced. As a SCR(6-7)-based FH-Fc construct, S2635 was shown in this study to have physiological ranges of binding affinity (K_D_ = ~10^−7^ M) selectively to CspA and CspZ from *B. burgdorferi* strain B31-5A4. In fact, CspA but not CspZ is produced in the spirochetes residing in feeding nymphs, which is required for bacteria to evade the complement in tick bloodmeal (24, 25). After *B. burgdorferi* invades a vertebrate host, CspZ but not CspA is produced to promote the systemic spread of bacteria (24, 26, 27). These findings thus support the possibility of S2635-mediated bacterial killing in the enzootic cycle by binding to CspA to decrease the levels of tick-to-host transmission of Lyme borreliae and to CspZ to reduce the extent of bacterial dissemination. Further, we found S2782 uniquely binding to documented SCR19-20 binders, OspE variants from *B. burgdorferi* strain B31-5A4. OspE variants are produced on the surface of *B. burgdorferi* residing in fed nymphs and vertebrate hosts (24). OspE-targeted antibodies were reported to not reduce the bacterial loads in fed nymphs but significantly decrease bacterial burdens in hosts after tick-to-host transmission (52). Such phenotypes are similar to our findings in the S2782-treated mice and therefore support the protective mechanisms underlying S2782 by binding to pathogens via OspE after Lyme borreliae transmission. Additionally, CspA and OspE are produced when Lyme disease bacteria are in ticks, but our results showed that CspA-targeted FH-Fc (i.e., S2635) but not the OspE targeted FH-Fc (i.e., S2782) eliminated bacteria in fed nymphs. One possibility to address this distinction could be the lower expression of levels of OspE, compared to those of CspA, when bacteria are in fed nymphs (53).

When the work was extended to different Lyme borreliae species or strains, we found S2635 showing bacterial killing ability to selected bacterial strains and species but S2782 displaying a broader ability to eradicate all tested species and strains. These results suggest the potential of developing S2782 as an anti-tickborne therapeutic with great breadth. As the binding activity of FH-Fc constructs to their FH-binding partners from pathogens is one of the determining factors for their efficacy, our findings raise the possibility that S2635 and S2782 differ in their capability to bind to different spirochete FH-binding protein variants. In fact, CspA shares approximately 40% sequence identity among different Lyme borreliae species, and OspE variants from different strains within the same species display greater than 85% sequence identity (54–56). Such protein polymorphism is consistent with documented CspA and OspE variant-to-variant different levels of human FH-binding activity (25, 57, 58). Although CspZ is highly conserved (~98%) among different spirochete strains within the same species and moderately conserved (~80%) among different Lyme borreliae species, CspZ variants also display variant-specific FH-binding activity (27, 59). Here, we found S2782 binding to all tested Lyme borreliae OspE variants at similar levels (K_D_ = 4 x 10^−7^ to 8 x 10^−8^ M). S2635 binds strongly to CspA and CspZ variants from *B. afzelii* and the CspA variant from *B. burgdorferi* 297 (K_D_ = ~10^−7^ M), but weakly to both CspA and CspZ variants from *B. bavariensis* (K_D_ = ~10^−6^ M). However, this FH-Fc does not bind to the CspZ variant from *B. burgdorferi* 297 and both CspA and CspZ variants from *B. garinii*. Therefore, these results suggest that the sequence differences between tested CspA and CspZ variants may impact S2635-binding activity, but the sequence variation of tested OspE variants here appears to not impact the S2782-binding activity. Further, such differences in the S2635- and S2782-binding affinities among the variants of spirochete FH-binding proteins are correlated with the extent of their ability to kill tested Lyme borreliae or strains. Our results suggest that S2782 has a broader spectrum as a Lyme borreliae anti-infective.

We have shown the ability of S2635 and S2782 to impact Lyme borreliae survivability *in vitro* in the presence of complement, but these results do not exclude the involvement of other mechanisms of action for these FH-Fc constructs *in vivo*. In fact, several publications indicate that *B. burgdorferi* can be cleared *in vivo* by both Fc-mediated phagocytosis or complement-dependent phagocytosis (i.e., opsonophagocytosis) (60–64). As future work, we plan to further investigate the role of phagocytic clearance in contributing to FH-Fc-mediated *B. burgdorferi* eradication. In this study, we tested the model of targeting host FH with the enhancement of classical pathway killing by Fc for the prevention of Lyme borreliae infection. Such a preventive strategy, known as pre-exposure prophylaxis, would require the agents to have a long half-life to be of practical use in humans. Overall, this study demonstrated the concept of targeting Lyme borreliae FH-binding proteins with the enhancement of classical pathway killing by Fc for the prophylaxis of Lyme disease. Such an innovative immunotherapeutic approach provides an alternative option for Lyme disease prevention, suggesting the possibility of developing a broad-spectrum preventive platform for multiple pathogens.

## MATERIALS AND METHODS

### Ethics Statement

All mouse experiments were performed in strict accordance with all provisions of the Animal Welfare Act, the Guide for the Care and Use of Laboratory Animals, and the PHS Policy on Humane Care and Use of Laboratory Animals. The protocol (Docket Number 22-451) was approved by the Institutional Animal Care and Use Committee of Wadsworth Center, New York State Department of Health. All efforts were made to minimize animal suffering.

### Mouse, ticks, bacterial strains, and human serum

Four-week-old, female C3H/HeN mice were purchased from Charles River (Wilmington, MA). BALB/c C3-deficient mice were from in-house breeding colonies (25) and *Ixodes scapularis* tick larvae were obtained from BEI Resources (Manassas, VA). *Escherichia coli* strain BL21(DE3), M15 or DH5α and their derivatives were grown at 37°C or other appropriate temperatures in Luria-Bertani broth or agar, supplemented with kanamycin (50µg/mL) or ampicillin (100µg/mL) (**Table S5).** *B. burgdorferi* strain B31-5A4 (**Table S5**) was grown at 33°C in BSK II complete medium (65). Cultures of *B. burgdorferi* B31-5A4 was tested with PCR to ensure a full plasmid profile before use (66, 67), the rest strains used in this study was kept within ten passages to prevent the potential plasmid loss. *A. tumefaciens* GV3101 (pMP90RK) containing the binary vector pTRAkc-P19, encoding the post-transcriptional silencing suppressor P19, and each of the FH-Fc constructs was used for transient expression using a *N. benthamiana* expression system as indicated (**Table S5**). Uninfected human serum (CompTech, Tyler TX) was confirmed as seronegative for Lyme disease infection as described (27).

### Expression and purification of FH/Fc fusion proteins in tobacco plants

Nucleotide sequences encoding human FH SCR6-7 (aa residues 321-443 (Genbank #: NP_000177)) and human FH SCR19-20 (aa residues 1048-1231 (Genbank #: NP_000177)), incorporating the D1119G mutation (68)), designed to employ optimal codon usage for expression in *Nicotiana benthamiana*, were synthesized by GENEWIZ (South Plainfield, NJ). Similarly, nucleotide sequences encoding human CH2-CH3 domains from IgG3 (aa residues 130-346 (Genbank #: CAA67886.1)) were also synthesized for optimal codon usage for expression in *Nicotiana benthamiana*, by GENEWIZ. In S2635, the SCR6-7 and the human CH2-CH3 domains from IgG3 were placed at N- and C-terminus, respectively, connected with a linker (GGGGSGGGGSGGGGSS), followed by the IgG1 hinge sequence (EPKSCDKTHTCPPCP) (**Fig. 1B, Text S1**). To generate S2782, a portion of the IgG1 hinge sequence (DKTHTCPPCP), followed by the human CH2-CH3 domains from IgG3 was placed at the N-terminus whereas the SCR19-20 was placed at the C-terminus. We did not add a flexible linker sequence in S2782. Note that in S2782, the C-terminal three residues of Fc, PGK, were replaced by the residues GQC, to facilitate construct stability and resistance to protease cleavage (**Fig. 1B**, **Text S2**) (35). These synthetic sequences of S2635 and S2782 were then placed downstream of the signal peptide of the murine mAb24 heavy-chain (lph) (**Fig. 1B**, **Text S1 and S2**) (69). The entire synthetic sequences were cloned into the plant binary expression vector pTRAkc (PMID: 17412974).

These recombinant proteins were then produced via transient expression by whole-plant vacuum infiltration of *N. benthamiana* ΔXT/FT using *A. tumefaciens* GV3101 and pMP90RK vector, as described previously (70–72). We then purified and concentrated S2635 and S2782 using Protein A-MabSelect SuRe or PrismA affinity columns (GE HealthCare) as described(73). Protein concentrations were quantified using a UV spectrophotometer for the absorption at 280 nm and extinction coefficients predicted from the mature amino acid sequences (excluding signal peptides).

### Cloning, transfection, expression and purification of CspA, CspZ, and OspE variants from Lyme borreliae

DNA encoding histidine or glutathione-S-transferase (GST) tagged CspA, CspZ, and OspE variants from different Lyme borreliae strains or species (**Table S2**) was used to express and purify these proteins using an *E. coli* expression system as described previously (21, 25–27, 42, 74, 75). Basically, the DNA encoding both N-terminal histidine- and GST-tagged proteins were synthetized by Synbio Technologies (Monmouth Junction, NJ), followed by subcloning into the pET28a (Millipore Sigma, Burlington, MA), pQE30Xa (Qiagen, Germantown MD), and pGEX4T2 vectors (Cytiva, Marlborough, MA), respectively, via BglII/BamHI restriction sites, using the service from Synbio Technologies (Monmouth Junction, NJ) (**Table. S2**). After transforming the pET28a- or pGEX4T2-associated plasmids into *E. coli* strain B21(DE3) or the pQE30Xa-associated plasmids into *E. coli* strain M15 as described (25), the protein was purified as in our previous work (25).

### Mouse infection

Flat *I. scapularis* nymphs carrying *B. burgdorferi* strain B31-5A4 were generated as described previously using BALB/c C3-deficient mice (25, 76). At 24-h prior to infection via placing nymphs on C3H/HeN mice, these mice were intramuscularly injected with PBS buffer (control) or 0.2 mg/kg of S2635 or S2782. The ticks were allowed to feed until repletion. At 21 dpf, the above-mentioned flat and replete ticks and mouse tissues were then used to determine bacterial burdens and the tibiotarsus joints were used to determine the severity of arthritis in the section “Quantification of spirochete burdens and histological analysis of arthritis.” Mouse sera were utilized to define the Lyme disease bacterial seropositivity as described in the section “ELISAs.”

### Quantification of spirochete burdens and histological analysis of arthritis

DNA was extracted from the indicated mouse tissues to determine bacterial burdens, using quantitative PCR analysis as described (77). Note that spirochete burdens were quantified based on the amplification of *recA* using forward (GTGGATCTATTGTATTAGATGAGGCTCTCG) and reverse (GCCAAAGTTCTGCAACATTAACACCTAAAG) primers. The number of *recA* copies was calculated by establishing a threshold cycle (Cq) standard curve of a known number of *recA* gene extracted from strain B31-5A4, and burdens were normalized to 100 ng of total DNA. For the ankles that were applied to histological analysis of Lyme disease-associated arthritis (**Fig. 2H**), the analysis was performed as described (77). Images were scored based on the severity of inflammation on a scale of 0 (no inflammation), 1 (mild inflammation with less than two small foci of infiltration), 2 (moderate inflammation with two or more foci of infiltration), or 3 (severe inflammation with focal and diffuse infiltration covering a large area).

### ELISAs

Seropositivity of the mice after infection with *B. burgdorferi* was determined by detecting the presence or absence of IgG recognizing C6 peptides, as described previously (78), as this methodology has been commonly used for human Lyme disease diagnosis (37). The maximum slope of optical density/minute of all the dilutions was multiplied by the respective dilution factor, and the greatest value was used as representative of anti-C6 IgG titers (arbitrary unit (A.U.)). Seropositive mice were defined as mice with sera yielding a value greater than the threshold, the mean plus three-fold standard deviation of IgG values derived from uninfected mice.

### Borreliacidal assays

The ability of S2635 or S2782 to kill different Lyme borreliae species or strains was determined as described with modifications (74, 77). Briefly, S2635 or S2782 were incubated, at different concentrations, with the Lyme borreliae. We then mixed the FH-Fc-Lyme borreliae with complement-preserved human serum (CompTech, Tyler TX) at a final concentration of 40% (20% for *B. garinii* strains ZQ1 and PBr, as that was the maximal concentration that did not result in bacterial killing in the absence of FH-Fc (**Fig. S1**)). The mixture was incubated at 33°C for 24 hours. Surviving (motile) spirochetes were quantified by direct counting using dark-field microscopy and expressed as the proportion of S2635- or S2782-treated to untreated Lyme borreliae (those exposed to complement-preserved human serum only). The concentration of S2635 or S2782 that killed 50% of spirochetes (EC_50_) was calculated using dose-response stimulation fitting in GraphPad Prism 9.3.1.

### Surface plasmon resonance (SPR) analyses

The interactions of recombinant CspA, CspZ, or OspE proteins with S2635 or S2782 were determined using a Biacore T200 (Cytiva), similar to the work in our previous studies (79). Basically, 10 micrograms of human S2635 or S2782 were conjugated to a Protein A chip (Cytiva). Quantitative SPR experiments were used to determine the binding kinetics of the CspA, CspZ, OspE variants that display FH-Fc binding activity. Basically, 10 µl of increasing concentrations (0.0625, 0.125, 0.25, 0.5, 1 µM, and/or 2 µM) of CspA, CspZ, or OspE proteins were injected into the control cell and the flow cell immobilized with S2635 or S2782 at 30 μl/min in PBS at 25°C. To obtain the kinetic parameters of the interaction, sonogram data were fitted by means of BIAevaluation software version 3.0 (GE Healthcare), using the one step biomolecular association reaction model (1:1 Langmuir model), resulting in optimum mathematical fit with the lowest Chi-square values.

### Statistical analyses

Significant differences were determined with a Kruskal-Wallis test with the two-stage step-up method of Benjamini, Krieger, and Yekutieli (80) and two-tailed Fisher test (for seropositivity)(81), using GraphPad Prism 9.3.1. A p-value < 0.05 was used to determine significance.

## ACKNOWLEDGEMENTS

The authors thank George Chaconas, Peter Kraiczy, Volker Fingerle, and John Leong for providing *B. burgdorferi* strain B31-5A4 and 297, *B. afzelii* strains VS461 and PKo, *B. bavariensis* strains PBi, *B. garinii* strain PBr and ZQ1, and CspA or CspZ-producing *E. coli* strains. They appreciate Sanjay Ram for valuable advice of the bactericidal assays used in this study. The authors thank the Wadsworth Animal Core for assistance with Animal Care and Steve Eyles of University of Massachusetts at Amherst Biophysical Characterization Core for SPR. The authors also thank to Tissue Culture & Media Core in Wadsworth Center and Tufts Microbiology Media Core to prepare the reagents and media, and Tufts Comparative Pathology and Genomics Shared Resource to generate the histopathology slides. This work was supported by NIH grant R41AI152954 (J.M., YT., K.W., Y.L.), R44AI152954 (C.M., YT., K.W., Y.L.) and R21AI144891 (J. M., Y.L.). The funders had no role in study design, data collection, interpretation, or the decision to submit the work for publication. The Authors, K.W. and Y.T., were employed by Planet Biotechnology. The rest of the authors declare no conflicts of interest.

