## Supplemental Figure Legends, Texts, and Tables for "Complement therapeutic Factor H-IgG proteins as pre-exposure prophylaxes against Lyme borreliae infections"

**SUPPLEMENTAL MATERIALS**

**Supplemental Figure legends**

**Figure S1. The results from bactericidal assays identifying the minimal percentage of human serum that does not impact *B. garinii* ZQ1 and PBr survival.** Indicated concentrations of the untreated or heat-inactivated human serum (serum percentage) were incubated with *B. garinii* (*Bg*) strains **(A)** ZQ1 and **(B)** PBr (5 × 10^5^ cells ml^-1^) for 24-h.. The surviving spirochetes were quantified from three fields of view for each sample using dark-field microscopy. The work was performed on three independent experiments. The survival percentage was derived from the proportion of serum-treated to the PBS-treated spirochetes*.* Shown is one representative experiment, and in that experiment, the data points are the mean ± SEM of the survival percentage from three replicates. Asterisks indicate the statistical significance (p < 0.05, Mann-Whitney test) of differences in percent survival between indicated spirochete groups and the spirochete group incubated with heat-inactivated human serum at the indicated concentration of serum.

**Supplemental TEXT**

**TEXT S1. The amino acid sequence of S2635.** The SCR6-7 of human FH and the CH2-CH3 domain of the Fc from human IgG3 are highlighted in blue and yellow, respectively. Between the FH and Fc regions, the linker and the hinge region of the human IgG1 are shown in purple and red, respectively.

tlkpcdypdikhgglyhenmrrpyfpvavgkyysyycdehfetpsgsywdhihctqdgwspavpclrkcyfpylengynqnygrkfvqgksidvachpgyalpkaqttvtcmengwsptprciggggsggggsggggssepkscdkthtcppcpapellggpsvflfppkpkdtlmisrtpevtcvvvdvshedpevqfkwyvdgvevhnaktkpreeqynstfrvvsvltvlhqdwlngkeykckvsnkalpapiektisktkgqprepqvytlppsreemtknqvsltclvkgfypsdiavewessgqpennynttppmldsdgsfflyskltvdksrwqqgnifscsvmhealhnhftqkslslspgk

**TEXT S2. The amino acid sequence of S2782.** The signal peptide is indicated by the underline. The hinge region of the human IgG1 and the CH2-CH3 domain of the Fc from human IgG3 is shown in red and yellow, respectively. The mutated three C-terminal residues, GQC, at the C-terminal Fc is bolded. The SCR19-20 of human FH is highlighted in green.

dkthtcppcpapellggpsvflfppkpkdtlmisrtpevtcvvvdvshedpevqfkwyvdgvevhnaktkpreeqyNstfrvvsvltvlhqdwlngkeykckvsnkalpapiektisktkgqprepqvytlppsreemtknqvsltclvkgfypsdiavewessgqpennyNttppmldsdgsfflyskltvdksrwqqgnifscsvmhealhnhftqkslsls**gqc**dstgkcgppppidngGitsfplsvyapassveyqcqnlyqlegnkritcrngqwseppkclhpcvisreimenynialrwtakqklysrtgesvefvckrgyrlssrshtlrttcwdgkleyptcakr

**SUPPLEMENTARY TABLES**

**Table S1. EC_50_ values of S2635 and S2782 to different species and strains of Lyme and relapsing fever borreliae.**

| **EC_50_ (µg/ml)^a^** | **Lyme**  **borreliae** | | | | | | | |
| --- | --- | --- | --- | --- | --- | --- | --- | --- |
|  | ***B. burgdorferi*** | | ***B. afzelii*** | | ***B. bavariensis*** | ***B. garinii*** | | |
|  | **B31-5A4** | **297** | **VS461** | **PKo** | **PBi** | | **ZQ1** | **PBr** |
| **S2635** | 0.17±0.02 | 0.23±0.01 | 0.16±0.01 | 0.22±0.13 | 11.24±4.31 | | n.k.^b^ | n.k. |
| **S2782** | 1.32±0.35 | 1.56±0.46 | 1.66±0.32 | 2.10±0.32 | 2.20±0.61 | | 1.70±0.08 | 1.80±0.58 |

^a^The concentration of the FH-Fc constructs that kill 50% of indicated Lyme borreliae strains in 40% of human serum except that the human serum incubated with *B. garinii* strains ZQ1 and PBr were 20%. Shown is the mean ± standard deviation of the EC_50_ values derived from three experiments (three replicates per experiment).

^b^n.k., No killing.

**Table S2. The binding affinity (K_D_) S2635 and S2782 to OspE variants from Lyme borreliae.**

| **The source of OspE variants** | **FH-Fc constructs** | **K_on_ (10^5^s^-1^M^-1^)** | **K_off_ (s^-1^)** | **K_D_ (µM)** |
| --- | --- | --- | --- | --- |
| ***B. burgdorferi* B31 (ErpA)** | **S2635** | n.b.^a^ | n.b. | n.b. |
|  | **S2782** | 8.12±0.62 | 0.15±0.008 | 0.19±0.004 |
| ***B. burgdorferi* B31 (ErpP)** | **S2635** | n.b. | n.b. | n.b. |
|  | **S2782** | 7.33±0.65 | 0.16±0.014 | 0.23±0.013 |
| ***B. burgdorferi* 297** | **S2635** | n.b. | n.b. | n.b. |
|  | **S2782** | 25.43±1.76 | 0.20±0.079 | 0.099±0.049 |
| ***B. afzelii* PKo** | **S2635** | n.b. | n.b. | n.b. |
|  | **S2782** | 21.73±1.09 | 0.14±0.034 | 0.076±0.029 |
| ***B. garinii* ZQ1** | **S2635** | n.b. | n.b. | n.b. |
|  | **S2782** | 6.86±0.18 | 0.29±0.10 | 0.43±0.087 |
| ***B. bavariensis* PBi** | **S2635** | n.b. | n.b. | n.b. |
|  | **S2782** | 18.70±5.88 | 0.15±0.008 | 0.11±0.058 |

^a^n.b., No binding

**Table S3. The binding affinity (K_D_) S2635 and S2782 to CspA variants from Lyme borreliae.**

| **The source of CspA variants** | **FH-Fc constructs** | **K_on_ (10^5^s^-1^M^-1^)** | **K_off_ (s^-1^)** | **K_D_ (µM)** |
| --- | --- | --- | --- | --- |
| ***B. burgdorferi* B31** | **S2635** | 2.12±0.26 | 0.04±0.028 | 0.22±0.08 |
|  | **S2782** | n.b.^a^ | n.b. | n.b. |
| ***B. burgdorferi* 297** | **S2635** | 2.18±0.14 | 0.02±0.007 | 0.11±0.015 |
|  | **S2782** | n.b. | n.b. | n.b. |
| ***B. afzelii* PKo** | **S2635** | 2.51±0.11 | 0.03±0.024 | 0.12±0.041 |
|  | **S2782** | n.b. | n.b. | n.b. |
| ***B. garinii* ZQ1** | **S2635** | n.b. | n.b. | n.b. |
|  | **S2782** | n.b. | n.b. | n.b. |
| ***B. bavariensis* PBi (Bga66)** | **S2635** | 0.66±0.01 | 0.07±0.018 | 1.10±0.593 |
|  | **S2782** | n.b. | n.b. | n.b. |
| ***B. bavariensis* PBi (Bga71)** | **S2635** | 0.40±0.09 | 0.04±0.004 | 1.12±0.418 |
|  | **S2782** | n.b. | n.b. | n.b. |

^a^n.b., No binding

**Table S4. The binding affinity (K_D_) S2635 and S2782 to CspZ variants from Lyme borreliae.**

| **The source of CspZ variants** | **FH-Fc constructs** | **K_on_ (10^5^s^-1^M^-1^)** | **K_off_ (s^-1^)** | **K_D_ (µM)** |
| --- | --- | --- | --- | --- |
| ***B. burgdorferi* B31** | **S2635** | 3.67±0.68 | 0.07±0.007 | 0.20±0.04 |
|  | **S2782** | n.b.^a^ | n.b. | n.b. |
| ***B. burgdorferi* 297** | **S2635** | n.b. | n.b. | n.b. |
|  | **S2782** | n.b. | n.b. | n.b. |
| ***B. afzelii* VS461** | **S2635** | 3.68±0.34 | 0.05±0.004 | 0.15±0.021 |
|  | **S2782** | n.b. | n.b. | n.b. |
| ***B. garinii* PBr** | **S2635** | n.b. | n.b. | n.b. |
|  | **S2782** | n.b. | n.b. | n.b. |
| ***B. bavariensis* PBi** | **S2635** | 1.61±0.11 | 0.15±0.01 | 0.98±0.306 |
|  | **S2782** | n.b. | n.b. | n.b. |

^a^n.b., No binding

**Table S5. Strains and plasmids used in this study.**

| **Strain or plasmid** | **Genotype or characteristic** | **Source** |
| --- | --- | --- |
| *B. burgdorferi* | | |
| B31-5A4 | Clone 5A4 of *B. burgdorferi* B31 isolated from *I. scapularis* ticks in US. | (83) |
| 297 | Clone A11/B11 of *B. burgdorferi* 297 isolated from human Cerebrospinal fluid of a Lyme disease patient in US. | (84, 85) |
| *B. afzelii* |  |  |
| VS461 | Clone JL of *B. afzelii* VS461 isolated from *I. ricinus* ticks in Switzerland. | (86) |
| PKo | Clonal isolate of *B. afzelii* PKo isolated from the skin lesion of erythema migrans from a Lyme disease patient in Germany. | (57) |
| *B. bavariensis* |  |  |
| PBi | Clonal isolate of *B. bavariensis* PBi isolated from human Cerebrospinal fluid of a Lyme disease patient from Germany. | (87) |
| *B. garinii* |  |  |
| ZQ1 | Clonal isolate of *B. garinii* ZQ1 isolated from *I. ricinus* ticks in Germany | (39) |
| PBr | Clonal isolate of *B. garinii* PBr isolated from Cerebrospinal fluid of a Lyme disease patient in Germany. | (88) |
| *E. coli* | | |
| DH5α | F- Φ80lacZΔM15 Δ(lacZYA-argF) U169 recA1 endA1 hsdR17(rk-, mk+) phoA supE44 thi-1 gyrA96 relA1 λ- | ThermoFisher |
| M15 [Prep4] | F-, Φ80ΔlacM15, thi, lac-, mtl-, recA+ , KanR^a^ | Qiagen |
| M15 [Prep4]/pQE30-CspA_B31_ | M15 producing histidine tagged residue 26 to 252 of CspA (BBA68) from *B. burgdorferi* strain B31 | (25) |
| M15 [Prep4]/pQE30-CspA_297_ | M15 producing histidine tagged residue 23 to 251 of CspA (AB1P08_04595) from *B. burgdorferi* strain 297 | This study |
| M15 [Prep4]/pQE30-CspA_PKo_ | M15 producing histidine tagged residue 28 to 242 of CspA (MMSA71) from *B. afzelii* strain PKo | (89) |
| M15 [Prep4]/pQE30-CspA_ZQ1_ | M15 producing histidine tagged residue 27 to 256 of CspA (ZQA68) from *B. garinii* strain ZQ1 | (89) |
| BL21(DE3) | F−, *ompT* *hsdSB* (rB− mB−) *gal dcm* (DE3) | Novagene |
| BL21(DE3)/pGEX4T2 | BL21(DE3) producing GST | (27) |
| BL21(DE3)/pGEX4T2-CspZ_B31_ | BL21(DE3) producing GST-tagged residues 20 to 230 of CspZ (BBH06) from *B. burgdorferi* strain B31 | (26) |
| BL21(DE3)/pGEX4T2-CspZ_297_ | BL21(DE3) producing GST-tagged residues 20 to 240 of CspZ (AB1P08_05365) from *B. burgdorferi* strain 297 | This study |
| BL21(DE3)/pGEX4T2-CspZ_VS461_ | BL21(DE3) producing GST-tagged residues 24 to 280 of *B. afzelii* strain VS461 | (77) |
| BL21(DE3)/pGEX4T2-CspZ_PBr_ | BL21(DE3) producing GST-tagged residues 21 to 280 of *B. garinii* strain PBr | This study |
| BL21(DE3)/pGEX4T2-ErpP_B31_ | BL21(DE3) producing GST-tagged residues 21 to 186 of ErpP (BBN38) from *B. burgdorferi* strain B31 | (90) |
| BL21(DE3)/pGEX4T2-ErpA_B31_ | BL21(DE3) producing GST-tagged residues 25 to 177 of ErpA (BBP38) from *B. burgdorferi* strain B31 | (75) |
| BL21(DE3)/pGEX4T2-OspE_PKo_ | BL21(DE3) producing GST-tagged residues 21 to 182 of OspE (BafPKo_N0036) from *B. afzelii* strain PKo | This study |
| BL21(DE3)/pGEX4T2-OspE_ZQ1_ | BL21(DE3) producing GST-tagged residues 1 to 158 of OspE (GenBank accession #: PQ337983) from *B. garinii* strain ZQ1 | This study |
| BL21(DE3)/pET28a-Bga66 | BL21(DE3) producing histidine tagged residues 23 to 250 of Bga66 from *B. bavariensis* strain PBi | This study |
| BL21(DE3)/pET28a-Bga71 | BL21(DE3) producing histidine tagged residues 23 to 251 of Bga71 from *B. bavariensis* strain PBi | This study |
| BL21(DE3)/pET28a-OspE_297_ | BL21(DE3) producing histidine-tagged residues 21 to 184 of OspE from *B. burgdorferi* strain 297 | (75) |
| BL21(DE3)/pET28a-CspZ_PBi_ | BL21(DE3) producing histidine tagged residue 21 to 280 of CspZ from *B. bavariensis* strain PBi | This study |
| BL21(DE3)/pET28a-OspE_PBi_ | BL21(DE3) producing histidine tagged 21 to 181 of OspE (GenBank accession #: PQ337984) from *B. bavariensis* strain PBi | This study |
| Plasmids |  |  |
| pQE30 | AmpR^b^; histidine tagged protein expression vector | Qiagen |
| pQE30-CspA_297_ | AmpR; pQE30 encoding histidine tagged residues 23 to 251 of CspA (AB1P08_04595) from *B. burgdorferi* strain 297 |  |
| pGEX4T2 | AmpR; GST-tagged protein expression vector | Qiagen |
| pGEX4T2-CspZ_297_ | AmpR; pGEX4T2 encoding GST-tagged residues 20 to 240 of CspZ (AB1P08_05365) from *B. burgdorferi* strain 297 | This study |
| pGEX4T2-CspZ_PBr_ | AmpR; pGEX4T2 encoding GST-tagged residues 21 to 280 of *B. garinii* strain PBr | This study |
| pGEX4T2-OspE_PKo_ | AmpR; pGEX4T2 encoding GST-tagged residues 21 to 182 of OspE (BafPKo_N0036) from *B. afzelii* strain PKo | This study |
| pGEX4T2-OspE_ZQ1_ | AmpR; pGEX4T2 encoding GST-tagged residues 1 to 158 of OspE (GenBank accession #: PQ337983) from *B. garinii* strain ZQ1 | This study |
| pET28a-Bga66 | KanR; pET28a encoding histidine tagged residues 23 to 250 of Bga66 from *B. bavariensis* strain PBi | This study |
| pET28a-Bga71 | KanR; pET28a encoding histidine tagged residues 23 to 251 of Bga71 from *B. bavariensis* strain PBi | This study |
| pET28a-CspZ_PBi_ | KanR; pET28a encoding histidine tagged residues 21 to 280 of CspZ from *B. bavariensis* strain PBi | This study |
| pET28a-OspE_PBi_ | KanR; pET28a encoding histidine tagged residues 21 to 181 of OspE (GenBank accession #: PQ337984) from *B. bavariensis* strain PBi | This study |

^a^ Kanamycin resistant

^b^ Ampicillin resistant
