## Supplementary figures and images for "Complement therapeutic Factor H-IgG proteins as pre-exposure prophylaxes against Lyme borreliae infections"

### Figure S1

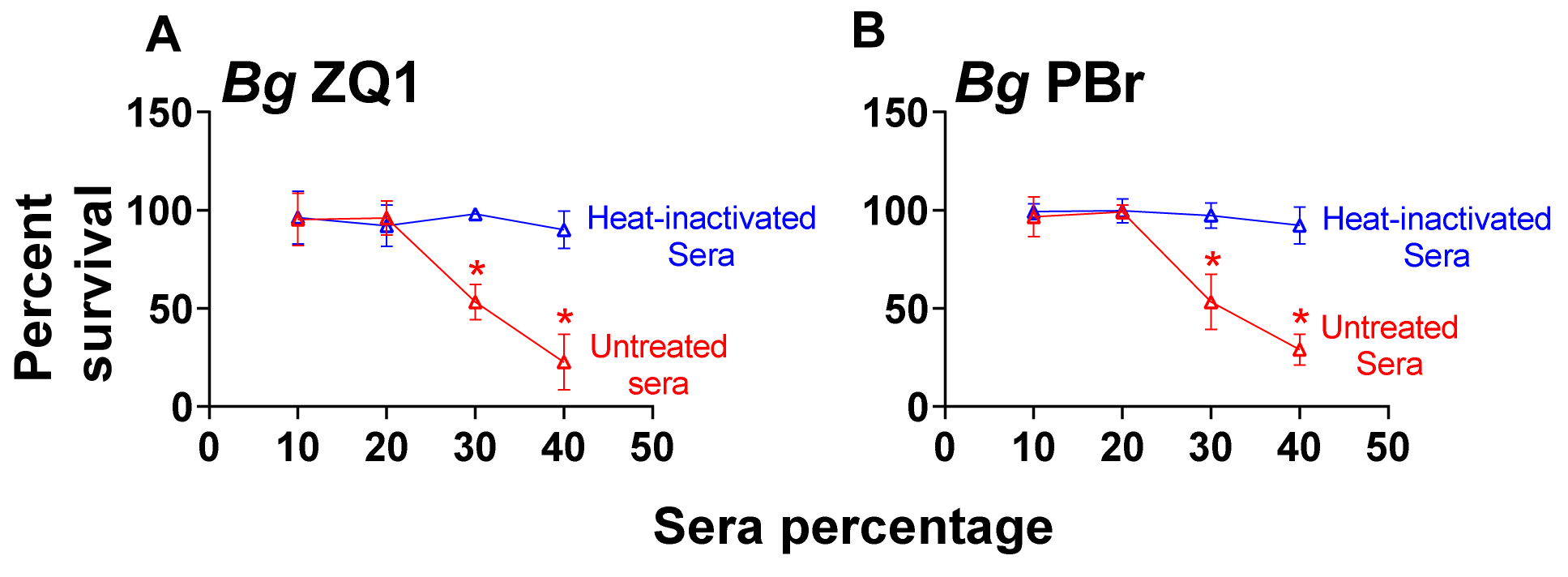
